# Single-cell RNA sequencing reveals dysregulation of spinal cord cell types in a severe spinal muscular atrophy mouse model

**DOI:** 10.1101/2022.05.30.493965

**Authors:** Junjie Sun, Jiaying Qiu, Qiongxia Yang, Qianqian Ju, Ruobing Qu, Xu Wang, Liucheng Wu, Lingyan Xing

## Abstract

Although spinal muscular atrophy (SMA) is a motor neuron disease caused by the loss of survival of motor neuron (SMN) proteins, there is growing evidence that non-neuronal cells play important roles in SMA pathogenesis. However, transcriptome alterations occurring at the single-cell level in SMA spinal cord remain unknown, preventing us from fully comprehending the role of specific cells. Here, we performed single-cell RNA sequencing of the spinal cord of a severe SMA mouse model, and identified ten cell types as well as their differentially expressed genes. Using CellChat, we found that cellular communication between different cell types in the spinal cord of SMA mice was significantly reduced. A dimensionality reduction analysis revealed 29 cell subtypes and their differentially expressed gene. A subpopulation of vascular fibroblasts showed the most significant change in the SMA spinal cord at the single-cell level. This subpopulation was drastically reduced, possibly causing vascular defects and resulting in widespread protein synthesis and energy metabolism reductions in SMA mice. This study reveals for the first time a single-cell atlas of the spinal cord of mice with severe SMA, and sheds new light on the pathogenesis of SMA.

## Introduction

Spinal muscular atrophy (SMA), a common lethal genetic disorder in children, is caused by mutations or deletions in the *survival of motor neuron 1* (*SMN1*) gene (MIM 600354), which leads to failure in encoding the SMN protein (UniProt accession number Q16637-1) [1]. Its pathology is characterized by motor neuron degeneration in the anterior horn of the spinal cord and necrosis of the neuromuscular junction [2]. Death in patients with SMA is due to respiratory failure caused by autonomic muscle denervation. Humans have a backup gene for *SMN1*, that is, *survival of motor neuron 2* (*SMN2*, MIM 601627), which can produce a small amount of functional SMN proteins due to mis-splicing but cannot compensate for *SMN1* loss [3]. Direct supplementation of SMN proteins or the correction of *SMN2* splicing is the main approach used for SMA treatment. In recent years, three effective drugs for SMA treatment have been approved by the US Food and Drug Administration, namely nusinersen, risdiplam, and abeparvovec-xioi. Nusinersen is an antisense oligonucleotide, and risdiplam is a small-molecule compound; these two drugs correct *SMN2* splicing; thus, increasing SMN protein levels. Abeparvovec-xioi is an adeno-associated viral vector-based gene therapy drug that increases SMN expression through the delivery of the full-length SMN coding sequence [4] [5, 6].

SMN is a functionally diverse housekeeping protein [7]. Its most well-understood function is to form a complex with the Gemin protein to participate in spliceosome assembly [8]. Another essential function of SMN is to participate in messenger RNA transport in motor neuron axons [9]. However, no SMN downstream molecules have been identified so far that causes motor neuron degeneration due to splicing errors or altered localization. SMN influences the SMA condition in a dose-dependent manner. In general, the more the copies of the *SMN2* gene, the milder the clinical phenotype of SMA. In addition, several modifiers affect the clinical phenotype of patients with SMA. Patients carrying the c.859G>C mutation on exon 7 and A-44G mutation on intron 6 of *SMN2* had a mild phenotype; the c.859G>C mutation creates an SF2/ASF-dependent exonic splice enhancer, and the A-44G mutation disrupts the HuR-dependent intronic splicing silencer [10, 11]. Furthermore, the overexpression of plastin 3 and neurocalcin delta can improve the clinical phenotype of patients with SMA [12, 13].

Numerous studies have proposed that in SMA, motor neurons undergo non-cell-autonomous death. Specific depletion of SMN in motor neurons causes limited dyskinesia rather than complete SMA [14]. Although a specific increase of SMN protein in motor neurons improves the pathological features and motor functions of SMA mice, the improvement in mouse lifespan is limited [15]. Glial cells are one of the most common cell types in the central nervous system (CNS), and their primary function is to maintain spinal cord homeostasis and nourish neurons. As a result, glial cells are the most likely to be involved in the non-cell-autonomous death of motor neurons. Astrocytes in SMA undergo morphological and functional changes that affect calcium homeostasis, microRNA formation, and protein secretion, reducing communication with motor neurons [16–19]. Surprisingly, SMN-specific supplementation in astrocytes did not prevent motor neuron death in SMA mice, but it did extend their lifespan [20]. In addition, other cell types in the spinal cord, such as vascular cells, are also disrupted in SMA [21]. Because the spinal cord is such a complex and heterogeneous tissue, different cell types or subtypes may play different roles in SMA pathogenesis. Therefore, an objective and comprehensive understanding of the transcriptome alterations in each cell type during the SMA process, as well as their effects on neurons, is critical.

Several studies have used transcriptomics to explore SMA pathogenesis, revealing a plethora of gene expression and splicing changes in SMA mice tissues or patient cells [22–28]. However, these studies have not identified which cell types in the SMA spinal cord are diseased or what transcriptome changes these cell types experience. Single-cell RNA sequencing (scRNA-seq) is an emerging technique that can be used to investigate cellular heterogeneity. This technique has uncovered disease-specific cell populations and altered transcripts that have never been reported before in Alzheimer’s disease (AD), amyotrophic lateral sclerosis (ALS), multiple sclerosis (MS), Parkinson’s disease (PD), Huntington’s disease (HD), and other diseases, providing new insights into these neurodegenerative diseases with unclear mechanisms [29–34].

The spinal cords of severe SMA mice and heterozygous mice were analyzed using scRNA-seq. A total of 22,155 cells were identified, and they were classified into 10 cell types, including oligodendrocyte precursor cells (OPCs), oligodendrocytes (OLs), committed OL precursors (COPs), neurons, schwann cells, blood cells, astrocytes, microglia, ependymal cells, and vasculature. We analyzed cell-type-specific differentially expressed genes (DEGs) and cell-cell communication between the different cell types. In addition, we identified cell subtypes and disease-specific genes for each subtype. Finally, we found that in SMA mice, fibroblasts were substantially reduced, emphasizing the importance of vascular defects in SMA pathogenesis.

## Results

### scRNA-seq and cell-type identification of the spinal cords of SMA and control mice

scRNA-seq was performed on Type I Taiwanese SMA mice at postnatal day 4 (P4). These mice exhibit severe SMA with a lifespan of approximately 10 days [35]. Studies have stated that treatment with nusinersen at 1-3 days postnatal instead of 5-7 days postnatal can improve the lifespan of these mice, suggesting that P4 is a turning point in the progression process of SMA pathology [36].

To explore disease-specific cell types and cell-type-specific DEGs, we first constructed a single-cell atlas. The spinal cords of three SMA mice (Smn^−/−^, SMN2^2tg/0^) at P4 were pooled as one sample for single-cell digestion, and heterozygous mice (Smn^+/–^, SMN2^2tg/0^) from the same litter were used as controls. Individual cells from both samples were subjected to scRNA-seq based on a 10× Genomics protocol. After quality-control filtering, 8887 control cells and 13,268 SMA cells were used for subsequent analysis, with an average median of 1933 genes per cell. To analyze global gene expression changes in the spinal cord, we constructed a bulk RNA sequencing (bulk-seq) atlas from the spinal cord tissues of SMA and heterozygous mice of the same age. Bulk-seq detected >20000 genes in total.

The uniform manifold approximation and projection (UMAP) method was used for the visualization and classification of all cells. The cells were divided into 16 clusters (**Fig. 1A**). Based on the expression of known CNS cell-type-specific genes [29], they were mapped into 10 cell types: astrocytes (cluster 0), microglia (clusters 1 and 12), OPC (clusters 2, 4, and 7), COP (cluster 3), OL (cluster 11), neuron (cluster 16), ependymal cells (clusters 11 and 15), vasculature (clusters 6, 13, and 9), blood cells (clusters 5 and 8), and Schwann cells (cluster 14) (**Fig. 1B** **sup 1A, B)**. OPCs were the most abundant, accounting for 22.97% of the total cells, whereas the proportion of neurons was the lowest, accounting for 0.46% of the total cells (**Fig. 1C**). No significant difference was observed in the overall cell proportion between SMA and control mice (**Fig. 1D, E**). The proportion of neurons and ependymal cells in SMA mice was slightly higher than that in control mice, and the proportion of blood cells in SMA mice was slightly lower than that in control mice (**Fig. 1E**). The number of neuronal cells (cluster 16) which expressed Meg3, Tubb3 and Snhg11 was too small to determine whether this was a motor neuron or not [37–41]. This approach for single cell suspension may result in neuronal loss.

**Figure 1.**
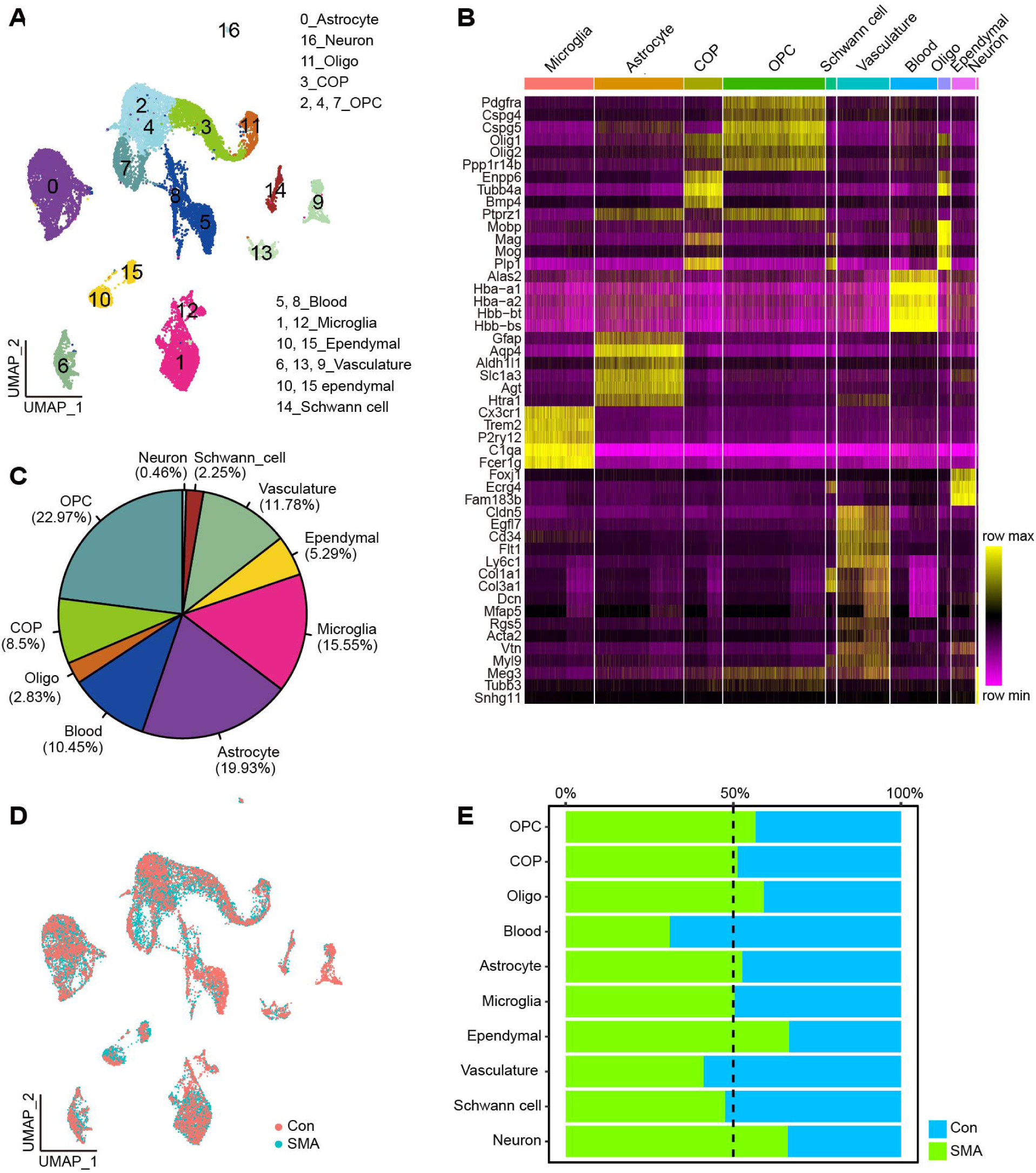
Cell type identification. (A) UMAP visualization showing the cluster of cell type from scRNA-seq. (B) The heatmap shows the average expression of marker genes for each cell type. The darker the yellow, the higher the expression level. (C) The proportion of each cell type to the total number of cells. (D) UMAP visualization showing the distribution of SMA and control cells. (E) Proportion of cells from SMA or control samples in each cell type.

### DEGs and functional analysis of SMA spinal cord cell types

We focused on cell type-specific DEGs in SMA and control mice. In most of the aforementioned cell types, approximately 5000–7000 genes were detected, with the exception of blood cells, where only 2008 genes were detected **(Sup 2A)**. According to log fold change (logFC) ≥ 0.2 and p < 0.01, the DEGs of each cell type were defined, with neurons, vasculatures, and COPs having the highest number of DEGs (960, 343, and 373, respectively) and OPCs, microglia, and ependymal cells having the lowest number of DEGs (45, 50, and 52, respectively) (**Fig. 2A** **and Sup File 1)**. Notably, in cell types other than blood cells, the number of downregulated genes was significantly higher than the number of upregulated genes, and this trend of predominantly downregulated expression was consistent with the results of bulk-seq (**Fig. 2B**).

**Figure 2.**
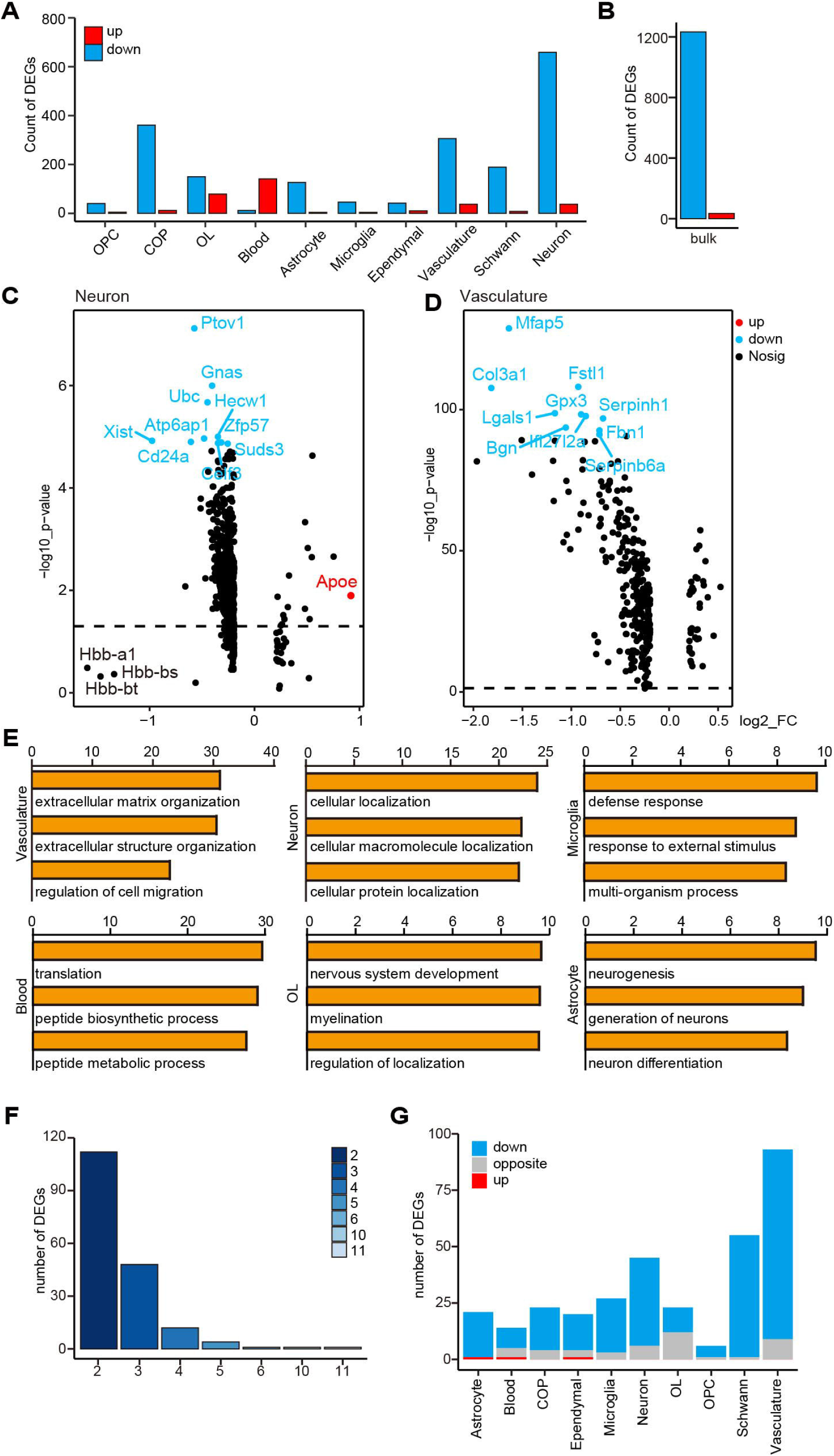
Analysis of DEGs and functions of scRNA-seq and bulk-seq. (A) The number of DEGs in different cell types in scRNA-seq. (B) Number of DEGs in bulk-seq. (C) A volcano plot showing DEGs in neurons. (D) Volcano plot showing DEGs of the vasculature. The x-axis represents logFC, and the y-axis represents log10 p-value. (E) GO analysis of DEGs in several cell types; the x-axis represents log10 p-value. Four cell types are not shown in the figure. The results of Schwann cells, OPCs, and COPs are similar to that of OLs, and the p-value of ependymal cells is >10^−5^. (F) An overlap gene analysis of cell-type-specific DEGs and bulk-seq DEGs; the y-axis represents the number of genes, and the x-axis represents the number of cell types; for example, the column with the column head “3” represents the number of overlapping DEGs of the bulk-seq and two cell types. (G) We analyzed whether the change trends of cell-type-specific DEGs in scRNA-seq and DEGs of bulk-seq are the same; blue indicates the same trend, and both are downregulated, red indicates the same trend, and both are upregulated, and gray indicates opposite trends.

Although neurons had the most DEGs, the fold change was small (**Fig. 2C**). Apoe, a microglial marker gene linked to neurodegenerative diseases like AD [42], was the most differentially upregulated gene. Hbb-a1, Hbb-bs, and Hbb-bt, which encode hemoglobin, were downregulated with the greatest fold in neurons, and their homolog Hbb-b1 was downregulated in another SMA mouse, Smn2B/− [43]. The X inactive specific transcript (Xist), a key regulator of mammalian X chromosome inactivation, was downregulated in most cells.

In vasculatures, a total of 15 genes were downregulated by more than two folds. From greatest to smallest fold changes, they were Timp1, Col3a1, Mfap5, Col1a2, Col1a1, Ccl7, Ccl2, Lgals1, S100a6, Ly6a, Bgn, Rarres2, Dcn, S100a10, and Cxcl1 (**Fig. 2D**). Ccl7 and Ccl2, for example, encode chemokines and act as downstream molecules of axonal SMN to mediate axonal growth [44]. Col3a1, Mfap5, and Dcn are all vascular fibroblast marker genes, implying that fibroblasts are the primary contributors to DEGs in SMA vasculature.

We next analyzed the biological processes (BP) of the DEGs in each cell type with the Gene Ontology (GO) database. DEGs in neuronal cells were mainly enriched in intracellular molecular localization; Schwann cells, astrocytes, and OL lineage cells were mainly enriched in nervous system development and neurogenesis; blood cells were enriched in protein synthesis; microglia were enriched in extracellular stimulus response and defense response; vasculature cells were enriched in altered extracellular components and cell migration (**Fig. 2E**). The majority of these non-neuronal cells’ GO terms were related to the cell membrane or extracellular matrix, implying that cellular communication in the SMA spinal cord is altered.

### Effects of SMN deficiency in different cell types are heterogeneous

We next performed an integrated analysis of data obtained from scRNA-seq and bulk-seq. After comparing DEGs in cell types with bulk-seq and counting the number of overlapping genes, we found that few DEGs appeared in three or more cell types, suggesting that SMN delficiency caused different gene expression changes in different cell types (**Fig. 2F**). In bulk-seq and scRNA-seq, only one DEG, Hbb-bt, was universally down-regulated, and another DEG, Hbb-a1, was downregulated in all cell types except COP **(Sup 2B)**. The most significant changes of Hbb-bt and Hbb-a1 were in the OL cells **(Sup 2C)**. Tmsb10 was the only gene that was upregulated in multiple cell types, with the highest expression in neurons and vasculature and with the most significant change in astrocytes **(Sup 2C)**. Most cell-type-specific DEGs were downregulated as shown in bulk-seq, and the vasculature matched the highest DEG numbers with bulk-seq (**Fig. 2G** **and Sup 2B)**.

The Kyoto Encyclopedia of Genes and Genomes (KEGG) pathway analysis revealed that most enriched pathways were only found in a small number of cell types or bulk-seq. Only two pathways, amino acid biosynthesis and phagocytosis, were significantly enriched in both bulk-seq and six cell types. Two pathways, fatty acid degradation and peroxisome functions, were significantly enriched only in bulk-seq **(Sup 2D)**. Though OPCs had a large number of DEGs, only the phagocytosis pathway was observed in the bulk-seq data. Notably, the differential pathways in the vasculature and microglia were the most similar to those found in bulk-seq, with overlaps in >50% of the pathways **(Sup 2D)**.

Our findings suggest that SMN deficiency had different effects on different cell types; bulk-seq was unable to adequately capture gene expression changes in single cell types; DEGs in the vasculature were the most similar to those found in bulk-seq.

### Cell–cell communications are reduced in the spinal cord of SMA mice

We used CellChat to analyze the communication network among spinal cord cells of control and SMA mice. CellChat offers two analysis methods based on the ligand–receptor pair: one uses secreted signaling interactions (SSIs) and the other uses cell–cell contact interactions (CCIs) [45]. Schwann and blood cells were removed because of their location in the peripheral nervous system and the low number of genes detected. The overall CCIs in the spinal cord of SMA mice were significantly reduced, with a 50% and 30% reduction in the number and strength of interactions, respectively (**Fig. 3A**). The cell types with the greatest reduction in CCIs, both in number and strength, were OPCs and COP cells, while microglia had the least significant reduction (**Fig. 3B, C**). The interactions between OPCs, COPs, and neurons were affected the most (**Fig. 3B, C**). In terms of the number and strength of interactions, SSIs in the spinal cord of SMA mice were reduced by 31% and 23%, respectively (**Fig. 3D**). Vasculature had the most significant reduction in the number of interactions, and OPCs and COPs had the most significant reduction in the strength of interactions (**Fig. 3E, F**). In SMA, the number and strength of SSIs decreased in most cell types, but increased in microglia (**Fig. 3E, F**), which could be related to the activation of microglia previously reported [46].

**Figure 3.**
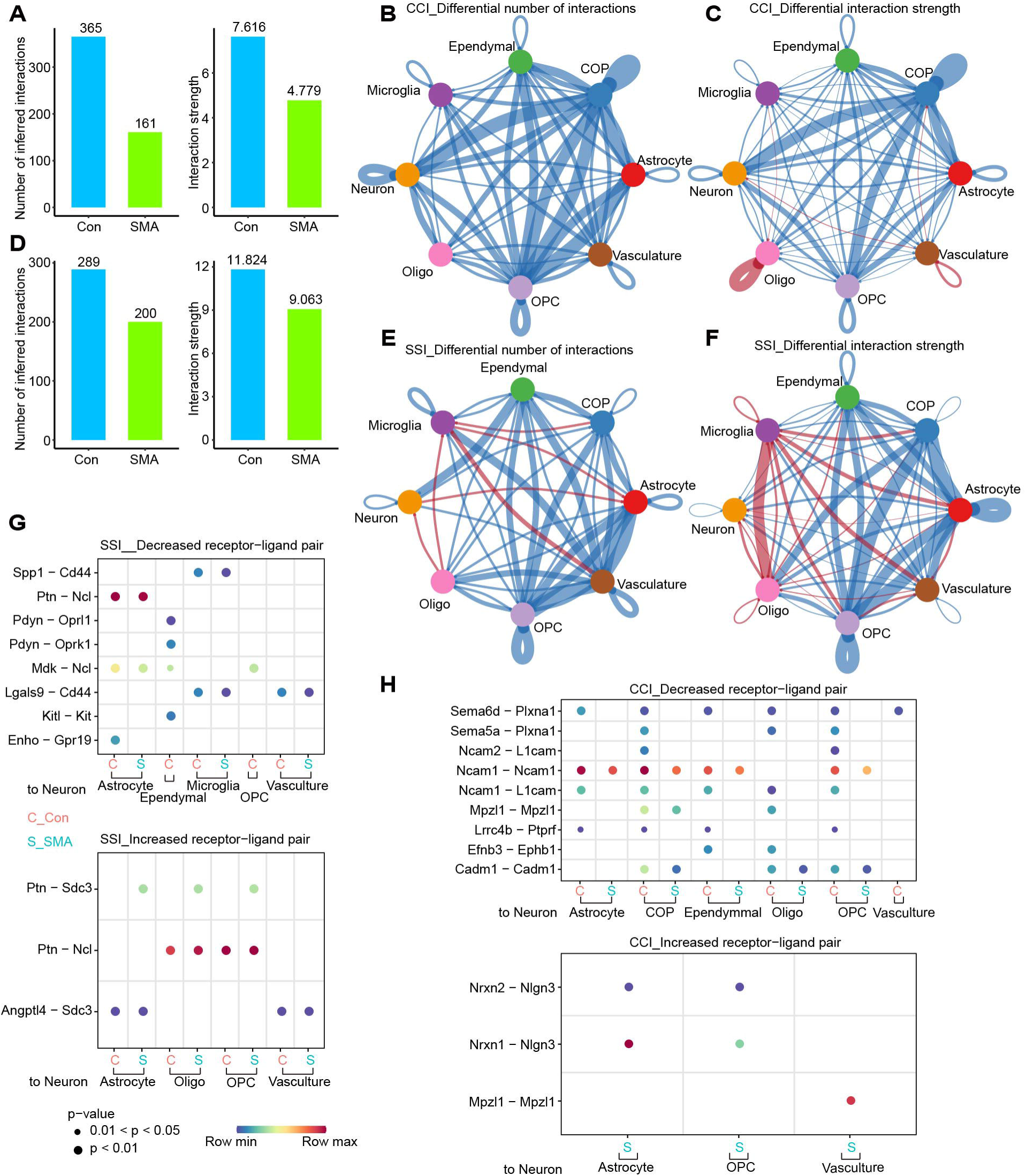
Cell-cell communications are reduced in the spinal cord of SMA mice. (A) Overall number and strength of cell-cell communications based on CCI. (B, C) Differences in the (B) number and (C) strength of CCIs communications of various cell types between SMA mice and control mice. The blue line indicates decreased communications, and the red line indicates increased communications. The thicker the line, the greater the difference. (D) Overall number and strength of cell-cell communications based on SSIs. (E, F) Differences in the number (E) and strength (F) of SSIs communications of various cell types between SMA and control mice. The blue line indicates decreased communications, and the red line indicates increased communications. The thicker the line, the greater the difference. (G, H) Receptor–ligand pairs that differ significantly between SMA and controls based on SSIs (G) and CCIs (H).

We were particularly interested in changes in neuronal communications with other cell types. In SMA, eight and three SSI-based pairings decreased and increased, respectively, and Ptn-Sdc3 was only detected in SMA and not in controls (**Fig. 3G**). SMA decreased in nine and increased in three CCI-based pairings, respectively (**Fig. 3H**). Sema–Plxna and Ncam–Licam pairs were only found in controls, whereas Nrxn–Nlgn3 and Mpzl1–Mpzl1 were only found in SMA (**Fig. 3H**).

### Identify cell subtypes and their DEGs

We next subdivided each cell type into subtypes. Astrocytes were divided into five clusters after dimensionality reduction (**Fig. 4A**). AS-2 had progenitor cell characteristics because its markers included cell cycle–related genes like Cdca8 and Stmn1 and were significantly enriched in the cell cycle pathway **(Sup 4A, B)**. AS-1 and AS-4 were defined as mature astrocytes because their marker genes were mostly found in the ribosome. The main difference between the two subtypes was that the main function of AS-1 was biased toward intracellular signal transduction, whereas AS-4 was more microglia-like, with high C1qa, C1qb, and Tyrobp expression **(Sup 3A, B)**. AS-3 was similar to a blood cell. Although AS-0 had the most cells, it was unable to be functionally defined due to a lack of signature genes.

**Figure 4.**
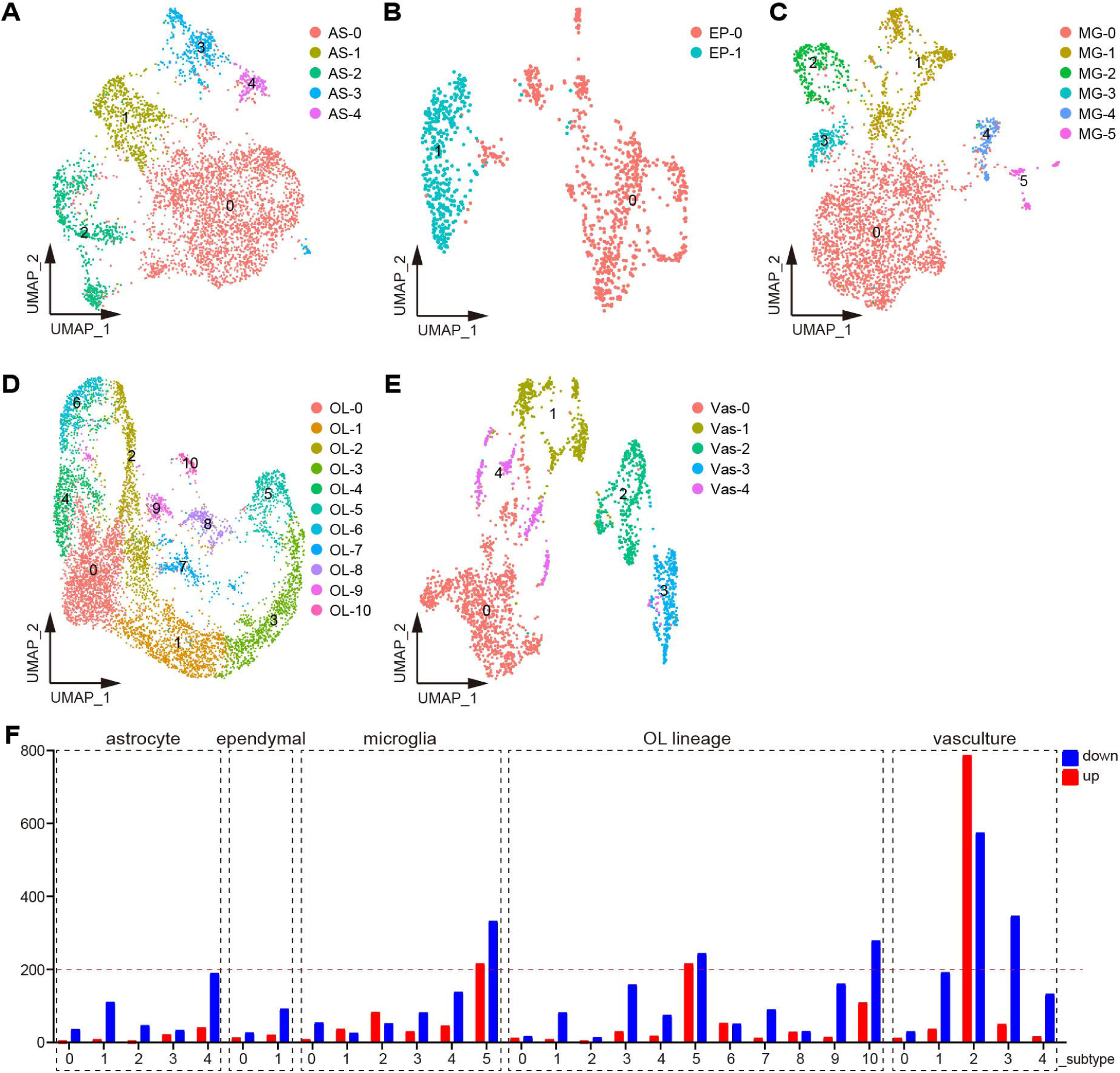
Identify cell subtypes and their DEGs. (A–E) Analysis of cell subtypes of (A) astrocytes, (B) ependymal cells, (C) microglia, (D) OL lineages, and (E) vasculature by UMAP. (F) Number of DEGs for each cell subtype.

Ependymal cells, which lines the central canal of the spinal cord, has a variety of functions. This population was dimensionally reduced into two clusters, with Ep-0 enriched for secretion and Ep-1 enriched for metabolism **(Fig. 4B, Sup 3C, D)**.

Microglia were classified into six subtypes (**Fig. 4C**). Of these, MG-0 was the most abundant, expressing most genes associated with microglial homeostasis, such as Cx3cr1, Trem2, and P2ry12, but it did not have specific markers compared to other subpopulations. Markers of the MG-1 subtype can be significantly enriched in the metabolic pathway, and thus MG-1 was defined as the metabolic microglia. The characteristic genes of MG-2 were Birc5, Cdca8, and Stmn1, and their functions were significantly enriched in the cell cycle and DNA replication pathways, and thus MG-2 was defined as dividing microglia. MG-3 cannot be functionally defined due to too few marker genes available. MG-4 specifically expressed Pf4, F13a1, and Lyve1, and these marker genes were enriched in the phagocytosis and endocytosis pathways, and thus MG-4 was defined as phagocytic microglia. MG-5 expressed the marker genes S100a11 and S100a6, which were significantly clustered in cell migration and cell adhesion, and was defined as migrating microglia **(Sup 3E, F)**.

Eleven OL lineage subtypes were identified (**Fig. 4D**). OL-5 was highly expressed with Mag, Mbp, and Mog, which were typical markers of myelinating OL. Markers in the axon guidance pathway were significantly enriched in OL-3, just as they were in OL5. However, because OL-3’s expression of myelin markers was low, it was classified as a nonmyelinating mature OL. OL-1 was specifically expressed with Bmp4 and Nkx2–2, which was defined as COPs. OL-2, OL-4, OL-6, and OL-10 were defined as OPCs because they were enriched with the cell cycle pathway, with OL-4 and OL-6 being OPCs with a robust dividing capacity. Although the cell cycle pathway was not enriched in OL-0, this subtype expressed OPC markers such as Pdgfra and Cspg5, and was therefore classified as OPCs **(Sup 3G, H)**.

Vasculatures contained five subtypes (**Fig. 4E**). Vas-0 and Vas-4 shared overlapping markers and had high levels of Cldn5, Pglyrp1, and Egfl7, all of which are markers for endothelial cells. The marker genes of Vas-1 were Rgs5 and Myl9, which were significantly enriched in the thermogenesis and oxidative phosphorylation pathways. Therefore, Vas-1 was defined as pericytes. Vas-2 and Vas-3, enriched with similar pathways, were defined as perivascular fibroblast subtypes as they both expressed Col3a1 and Dcn. The difference was that Vas-2 markers were significantly enriched with inflammation-related pathways such as interleukin-17 and tumor necrosis factor, whereas Vas-3 markers were not **(Sup 3I, J)**.

We compared the DEGs for each cell subtype in SMA and control mice. Vas-2 had 1362 DEGs, which is significantly higher than other subtypes, implying that Vas-2 is the most severely defective cell subtype in SMA mice’s spinal cord (**Fig. 4F**). MG-5, OL-5, OL-10, and Vas-3 also contained numerous DEGs, with 549, 460, 388, and 397, respectively **(Fig. 4F, Sup File 2)**.

### Altered gene expression of glial cell subtypes in the SMA spinal cord

We started by looking at changes in the numbers of major glial cells. No significant differences in cell numbers were observed in the subpopulations of the OL lineage and astrocytes in SMA compared to controls **(Sup 4A, B)**. However, the cell number of the migrating microglia MG-5 was reduced by approximately two-third in SMA mice compared with control mice **(Sup 4C)**. Although MG-5 has the most DEGs among the microglia subpopulation, neither the fold nor the p-value of the DEG changes were significant (**Fig. 5A**). Functional analysis revealed that downregulated genes in MG-5 in SMA were mainly associated with immune system response, translation, and RNA splicing, while upregulated genes were not functionally significant (**Fig. 5B**).

**Figure 5.**
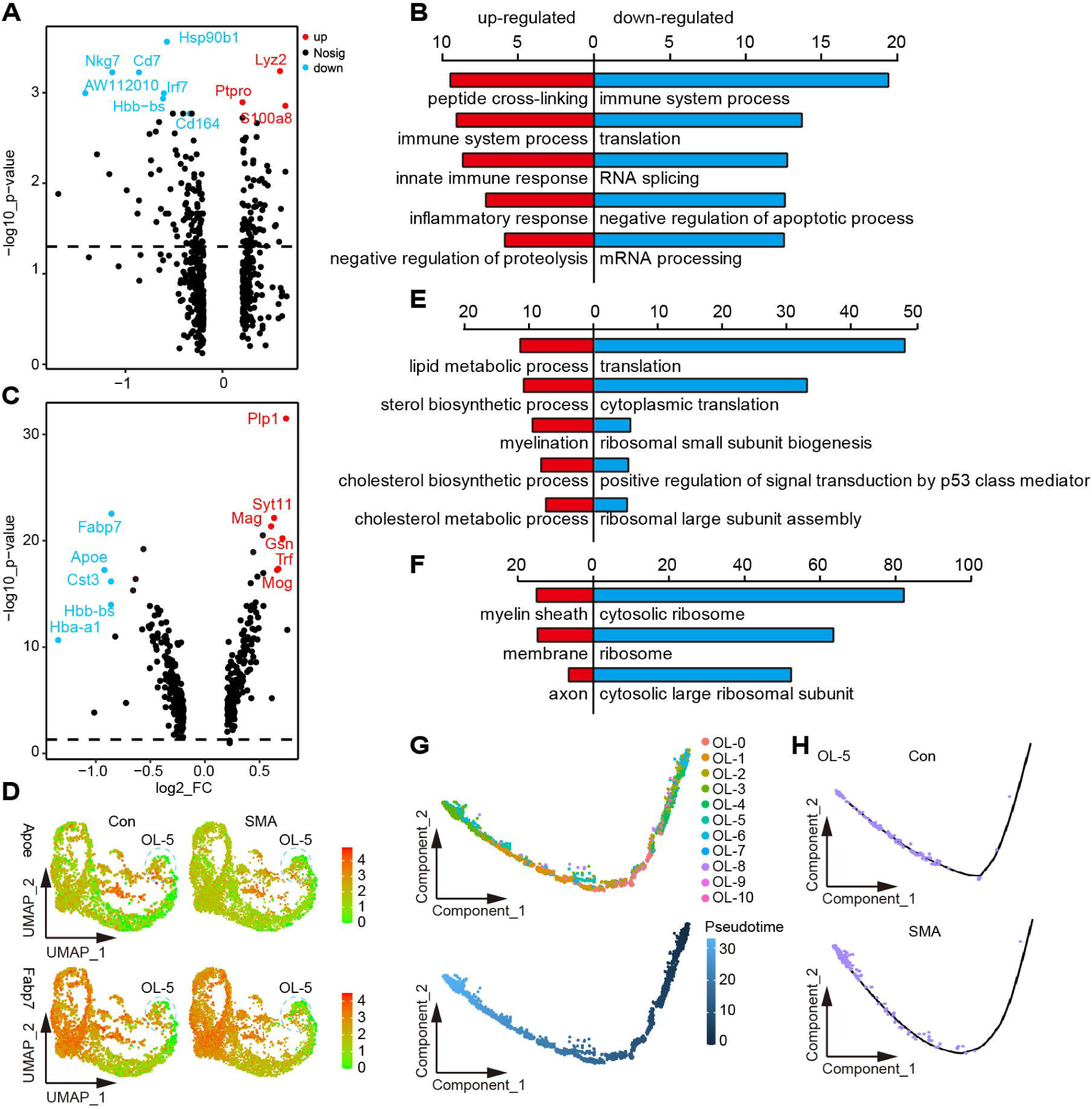
Altered gene expression of glial cell subtypes in the SMA spinal cord. (A) Volcano plot of DEGs in MG-5 subpopulations. (B) BP analysis of DEGs in MG-5 subtypes; the top 5 most significant terms are shown: red indicates upregulated terms, and blue indicates downregulated terms. (C) Volcano plot of DEGs in OL-5 subpopulations. (D) Signature DEGs expression of OL-5. (E, F) BP and cellular component analysis of DEGs in OL-5; the top 5 most significant terms are shown. (G) The position of each subtype of OL lineage on the pseudo time trajectory maturing from right to left. (H) Pseudo time trajectory of OL-5 in SMA (right) and control (left) mice.

OL-5 was the OL lineage subtype with the most DEGs, the most significant of which were Apoe and Fabp7 (**Fig. 5C, D**). Apoe encodes a member of the apolipoprotein A1/A4/E family, and its polymorphism is the strongest genetic risk factor for AD [47]. Fabp7 encodes a member of the fatty acid-binding protein family, which can bind to fatty acids, a crucial component of myelin [48, 49]. The BP analysis of DEGs revealed that downregulated DEGs were mainly related to protein synthesis, whereas upregulated DEGs were mainly enriched in myelination-related pathways (**Fig. 5E**). Consistently, the cellular component analysis of DEGs revealed that downregulated and upregulated genes were mainly localized to the ribosome and myelin sheath, respectively (**Fig. 5F**).

We performed a pseudotime analysis of OL lineage in SMA and control mice (**Fig. 5G**) and found that the OL-5 of SMA was more mature in the trajectory than that of controls, which was consistent with the results of the BP analysis (**Fig 5E, H**). Similarly, OL-0, OL-2, OL-4 and OL-10 cells appeared to be more mature, whereas OL-1 and OL-3 appeared less mature in SMA in the trajectory **(Sup 4D)**. Our results suggested that the maturation states of OL lineage cells may be affected in SMA mice.

### Subpopulation of perivascular fibroblasts was drastically reduced in SMA spinal cords

The Vas-2 subtype in SMA was drastically reduced (**Fig. 6A**). Both Vas-2 and Vas-3 were fibroblast subpopulations, and SMA selectively reduced the number of Vas-2 while leaving Vas-3 cells alone (**Fig. 6A, B**). Most fibroblast markers were downregulated in Vas-2, such as Col3a1, Col1a1, Dcn, and Fn1 (**Fig. 6C**). The gene with the greatest fold differential expression was Col3a1, which was downregulated nearly 4-fold (**Fig. 6C, D**). In addition, Rarres2, the gene encoding the angiogenesis regulator Chemerin, was downregulated by >2-fold in this cell subpopulation [50] (**Fig. 6C, E**). Notably, these DEGs that were significantly altered in both Vas-2 and Vas-3 fibroblast subtypes overlapped slightly (**Fig. 6F**), again indicating that the effects of SMN deficiency were cell-type specific.

**Fig. 6.**
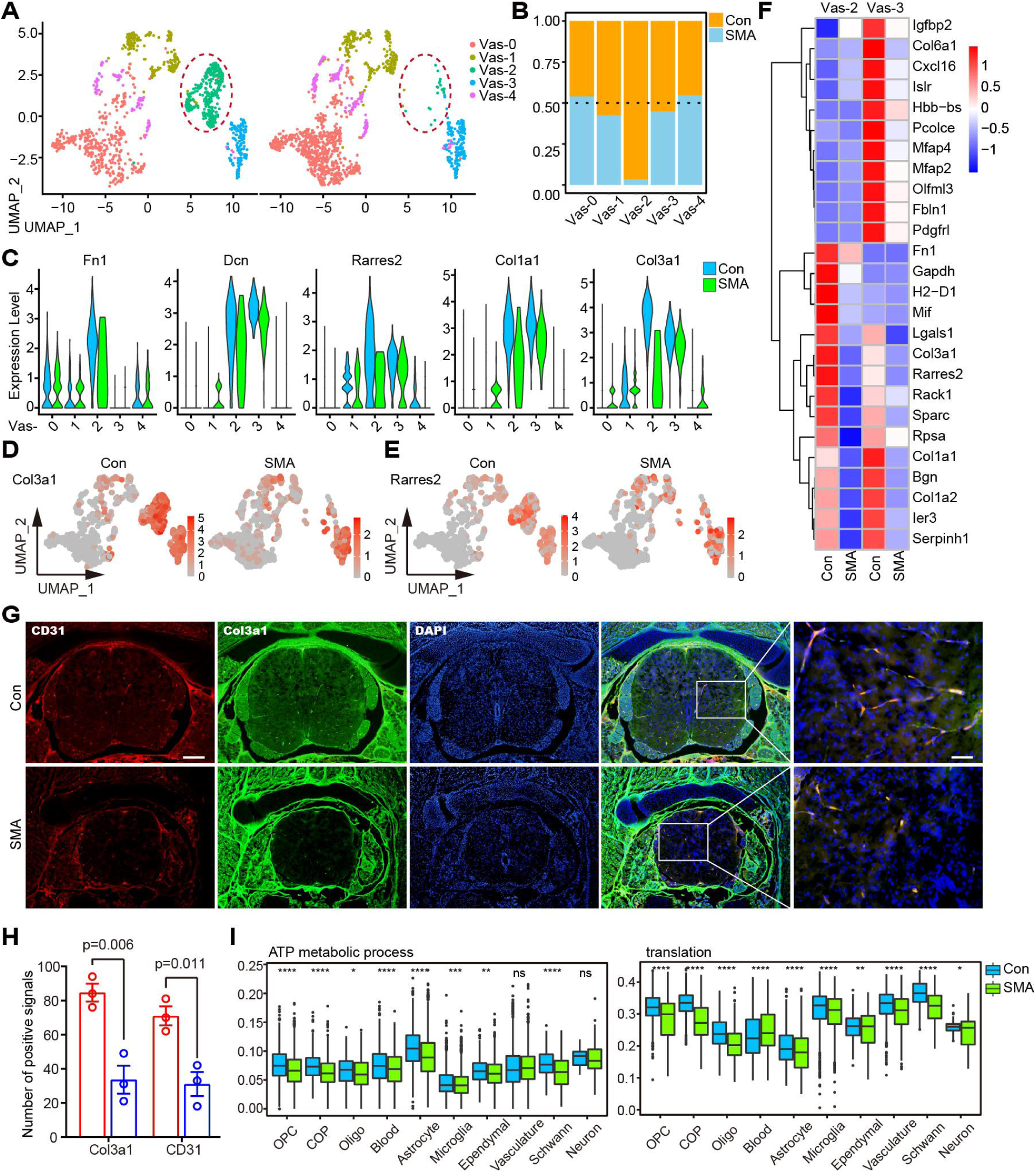
Subpopulation of perivascular fibroblasts was drastically reduced in SMA spinal cords. (A) UMAP visualization showing the subtypes of vasculature in control and SMA mice. Vas-2 is marked by red circles. (B) Comparison of the cell numbers of various subtypes in the vasculature of SMA and control mice. (C) Violin plots showing the expression of fibroblast marker genes in vasculature subtypes. (D, E) UMAP visualization showing the expression of Col3a1 and Rarres2 in the vasculature of SMA and control mice. (F) Expression analysis of the top 13 most significant DEGs in two fibroblast subtypes, Vas-2 and Vas-3. (G) Immunofluorescence staining of spinal cord transections in SMA and control mice. Here, 4’,6-diamidino-2-phenylindole (DAPI)-labeled nuclei are indicated in blue, Col3a1-labeled fibroblasts are indicated in green, and CD31-labeled vascular endothelial cells are indicated in red. Bar = 200µm (left) and 50 µm (right). (H) Statistical results of staining for CD31 and Col3a1. The P-value is calculated by Student’s t-test, n = 3, bars represent the mean with SEM. (I) Expression analysis of protein synthesis (translation) and energy metabolism (ATP metabolic process) pathways of various cell types in SMA and control mice. p-values were calculated through the t-test, ns: p > 0.05, *: p < 0.05, **: p < 0.01, ***: p < 0.001, ****: p < 0.0001.

We performed immunofluorescence experiments to identify the changes in the vasculature cells. Pecam1/CD31 and Col3a1 were blood vessel and fibroblast markers, respectively [21]. CD31 and Col3a1 were nearly 100% co-localized (**Fig. 6G**), indicating that vascular endothelial cells and fibroblasts coexisted in a spatial and functionally relevant manner, consistent with fibroblasts’ potential role as vessel supporting cell [51, 52]. Notably, along with the reduction of CD31 in the spinal cord of SMA at P4 [21], the expression of Col3a1 was dramatically decreased (**Fig. 6G, H**). When combined with our scRNA-seq data, this suggests that Vas-2 deficiency as a supporting cell may play a role in the vascular defects seen in SMA mice [21, 51, 52]. Consistently, we observed a downregulation in the energy metabolism or protein synthesis pathways across all cell types and the entire tissue **(Fig. 6I, Sup 5)**. This could we be due to the hypoxic environment caused by vessel defects [21].

## Discussion

Although motor neuron degeneration is a hallmark feature of SMA, increasing evidence suggests that SMA is a systemic disease caused by SMN protein deficiency [53]. Hua et al. confirmed that SMA is a non-cell-autonomous defect of motor neurons caused by non-motor neuron cytopathies [54]. However, the cell types in the spinal cord affected by SMA and the extent of involvement of the SMN protein in the different cell types are unexplored. Here, we used scRNA-seq and bulk-seq to analyze the genes and cell subpopulations affected in the spinal cords of severe SMA mice on P4 and attempted to fill the aforementioned knowledge gap.

In particular, we defined 10 cell types, including OPCs, OLs, COPs, neurons, Schwann cells, blood cells, astrocytes, microglia, ependymal cells, and vasculatures, and analyzed their DEGs caused in SMA. Compared with the bulk-seq data reported previously, this study is an advance in accurately identifying gene expression alterations in each cell type in the SMA spinal cord [22, 55–59]. Several studies have used laser capture microdissection to isolate motor neurons and glial cells, which greatly improved the purity of individual cell types, but this still did not completely rule out interference from other cell types [23, 25, 43, 60]. DEGs differ across cell types, suggesting the specific cellular responses of SMN protein deletion.

The downregulation of hemoglobin genes (Hba-a1 and Hbb-bt) was observed in most cell types and bulk-seq. Hemoglobin is known for binding oxygen in red blood cells, but they are also present in neurons and glial cells [61, 62]. Studies have confirmed that the downregulation of hemoglobin gene expression is associated with neurodegenerative diseases, such as AD and MS [63, 64]. In the spinal cord of SMAΔ7 mice, the hemoglobin-encoding genes Hbb-b2 and Hbb-b1 were downregulated by 2.6-fold at 7 days after birth [56]. Significant recovery of hemoglobin subunit alpha was observed in the cerebrospinal fluid of SMA patients treated with nusinersen [65]. Hemoglobin-encoding genes were not significantly altered in SMN knockout Hela cells, suggesting that they are not direct downstream molecules of SMN and may be the result of reduced cellular oxygen metabolism [22].

The bulk-seq and microarray data of type I Taiwanese SMA mice spinal cord showed that only a few genes were altered in presymptomatic mice (P1), whereas numerous gene expression changes were observed in the late stage of the disease (P5) [22, 57]. P4 is considered an intermediate state in disease progression and has received less attention before. In the P4 spinal cord of SMA mice, both scRNA-seq and bulk-seq revealed that the number of downregulated genes was much greater than the number of upregulated genes, a pattern that differed from that of P5. The cell division pathway and axon guidance genes were significantly downregulated in the P5 spinal cord, which was not observed in the P4, indicating that SMA pathogenesis undergoes great physiopathological changes from the middle stage to the late stage [22]. Most of the down-regulated pathways in the P4 spinal cord are metabolically related, consistent with the widespread metabolic dysregulation observed in SMA that has been demonstrated [66].

Normal communication between the various cell types of the spinal cord is required for the maintenance of neuronal structure and function, particularly during development [67]. Dysregulation of intercellular communication may contribute to SMA [68]. CellChat analysis revealed that intercellular communication in the SMA mice’s spinal cord was reduced in both number and intensity. Astrocytes are one of the most altered cell types. In SMA, a study using *in vitro* culture systems revealed abnormal interactions between astrocytes and motor neurons in SMA [19]. Furthermore, significant changes in communication between the cell types OPC and COP have been observed, but no study has yet confirmed whether this has an impact on SMA. Several ligand–receptor pairs were found to be differentially expressed in SMA and control mice. Ncam and L1cam are shown downregulated in SMA patients or motor neuron disease cell models, consistent with our results [69–71]. Although it is unclear whether these ligand-receptor pairs play a role in SMA pathogenesis, they still provide potential topics for follow-up research.

Numerous studies have focused on transcriptome alterations in SMA motor neurons. In addition to the aforementioned laser capture microdissection, several studies have analyzed the transcriptome of motor neurons produced from SMA patient-derived induced pluripotent stem cells. Although both Giacomo P Comi’s and Lee L Rubin’s teams obtained high-purity Hb9 positive cells, the DEGs and pathways discovered in the two studies were not identical due to differences in cell sources and different induction techniques [26–28]. There are numerous subtypes of motor neurons [72]. Ling et al. found that motor neurons do not innervate all muscles equally in mice with severe SMA in the late stages of the disease, but selectively denervate some of them (e.g., axial and appendicular muscles) [73]. Another study revealed that SMNs have different roles in maintaining the integrity of central and peripheral synapses [74]. Therefore, different neuron types may play different roles in SMA. However, only a small number (0.46%) of neuronal cells were identified in this study, which is insufficient to determine whether they are motor neurons and which subtype they belong to, which is a limitation of the study. This may be a result of the cell suspension technique we used for single cell isolate, in which neurons are more susceptible to death [75]. In the future, scRNA-seq of neurons sorted through flow cytometry can address this knowledge gap.

Microglia are immunocompetent cells of the CNS [76]. SMAΔ7 mice showed significant microglia activation in the late stages (after 10 days after birth) but not in the early and middle stages (before 7 days after birth) [77] [46, 78]. Aleksandra Vukojicic et al. reported that microglia leads to abnormal spinal cord motor circuits in SMAΔ7 mice by activating the canonical complement pathway [79]. However, this phenomenon was not evident in the early stage of the disease (P4), and pharmacological inhibition of the complement pathway had limited lifespan improvement in SMA mice. Therefore, microglial activation is a consequence rather than a cause of SMA. No reports are available regarding microglia activation in the spinal cord of type I Taiwanese mice. Our data classified microglia into six subtypes, with MG-5 migrating microglia being the most dissimilar between SMA-affected and control mice, suggesting that it is the subtype affected by SMA.

Astrocytes are one of the most abundant cell types in the CNS and play crucial roles in physiological processes such as maintaining ion homeostasis, nourishing and protecting neurons, and synapse development and plasticity [80]. Supplementation of SMN specifically to astrocytes doubles the lifespan of Δ7 SMA mice, suggesting that astrocytes can affect the severity of SMA [20]. Our scRNA-seq data revealed that the AS-4 reactive astrocyte subtype, which had higher “ribosomal” activity, was significantly different between SMA and control mice. The marker genes of AS-4 were C1qa, C1qb, and Tyrobp, and these genes had microglia-like characteristics. A subpopulation of reactive astrocytes known as A1 has been found to have highly upregulated expression of many classical complement cascade genes, similar to AS-4 found in this study [81, 82]. A1 loses its ability to maintain synaptic function, loses its ability to phagocytize, and is highly neurotoxic [81]. This subtype is associated with a variety of neurodegenerative diseases, including HD, AD, PD, MS, and ASL, and is triggered by normal aging [81, 83]. Our scRNA-seq data suggest that this subtype is also present in SMA and may play a role in SMA.

The traditional view is that OLs are myelination cells in the CNS. Many new functions of OL lineages beyond myelination have been gradually elucidated, such as providing metabolic support to neurons and regulating water and ion homeostasis [84]. Whether OL lineages are involved in SMA pathogenesis is controversial. O’Meara et al. found that OL growth, migration, differentiation, and myelination were not affected in severe SMA mice (Smn^−/−^, SMN2^+/+^) [85]. However, Kazuki Ohuchi et al. found that the OL lineage was impaired in Δ7 SMA mice [86]. Our data show that the subtype of OL lineages that is most affected by SMA is the myelinating OL subtype, OL-5. In this suntype, myelination signaling pathway is slightly up-regulated and the protein synthesis pathway was significantly reduced. Furthermore, the aforementioned COP cells and OPCs are the cell types in SMA that have the greatest reduction in cellular communication. Therefore, it would be fascinating to dissect out the specific changes in OL lineage cells in SMA.

The vasculature is a vital source of material and energy metabolism in the CNS, and the integrity of the vasculature is essential for the normal functioning of the spinal cord. Structural and functional abnormalities of the vasculature are common in neurodegenerative diseases [87, 88]. Vascular defects have also been identified in SMA patients and mouse models [21, 89–91]. Two fibroblast subpopulations were significantly altered in SMA, with one subpopulation experiencing a precipitous decrease in number. Our findings refine the affected vascular subpopulations, which have crucial implications for exploring the cellular targets of SMA. Fibroblasts are a group of cells that attach to the outer walls of blood vessels and are found in all blood vessels except capillaries [51, 92, 93]. The well-known function of fibroblasts is to participate in fibrotic scar formation after spinal cord injury [94]. The fibroblast markers Col3a1 and Dcn are greatly upregulated after spinal cord injury [95]. In addition, fibroblasts transmit signals through the bloodstream, affecting the progression of neurodegenerative diseases. ALS patients in whom fibroblast protein product accumulates in perivascular space have shorter survival times [96].

In spite of the fact that fibroblasts are an important component of the vasculature, there hasn’t been much research into their precise function [88]. The roles of fibroblast have only just recently begun to attract attention again due to the discovery of its inefficiency in neurodegenerative disorders by scRNA-seq. Andrew C Yang et al. pointed out that fibroblasts may be involved in ion and amino acid transport in the CNS [97]. Our research is the first of its kind to report on the fibroblast defects that are associated with SMA. It is important to conduct research to determine whether or not fibroblast defects are typical of neurodegenerative diseases. It’s also fascinating to learn more about fibroblast functions other than supporting cells, as well as why it’s more vulnerable in neurodegenerative disorders.

We observed widespread inhibition of energy metabolism and protein synthesis pathways across cell subtypes. Alterations in these pathways may be another manifestation of morphological and functional defects in mitochondria in SMA mouse motor neurons [98]. Somers et al. demonstrated that the spinal cord of SMA mice has a defect in the blood–spinal cord barrier with significant functional hypoxia [21]. The weakened interaction between vascular and CNS cells in our findings also suggests that an abnormal blood vessel structure may lead to decreased material exchange with CNS cells, affecting spinal cord function to some extent.

In summary, this study provides the first single-cell atlas of SMA. Through scRNA-seq, we identified the disease-specific DEGs of each cell type and subtypes of SMA spinal cord and found that the decline in the number of vascular fibroblasts was most pronounced at the single-cell level. These previously unreported finding shed new light on the pathogenesis of SMA.

## Materials and Methods

### Animal management

The experimental animals in this study were housed in the Specific Pathogen Free barrier provided by the Animal Center of Nantong University under the following conditions: temperature (22°C ± 2°C), humidity (60% ± 5%), and light of Alternate dark/light for 12 hours. Animal experiments were approved by the Animal Ethics Committee of Nantong University (No.2150409) and followed the International guidelines (NIH) for the Care and Use of Laboratory Animals.

### Generation of transgenic mice

The Taiwanese SMA mice used in this study was a gift from Professor Hua Yimin at Soochow University and was originally purchased from the Jackson Laboratory (FVB.Cg-Smn1tm1HungTg (SMN2)2Hung/J, stock number 005058) [35, 99]. There are two genotypes of this mouse, severe (Smn^-/-^, SMN2^2tg/0^) and mild (Smn^-/-^, SMN2^2tg/2tg^). Severe and control mice (Smn^+/-^, SMN2^2tg/0^) were sequenced in this project, generated by crossing Het knockout mice (Smn^+/-^) with mild Taiwanese mice (Smn^-/-^, SMN2^2tg/2tg^). Genotyping of tail-end tissue from newborn mice was performed using the method described in One step mouse genotyping kit (PD101-01, vazyme, nanjing, china). A Primer pair (F: 5’-AGCCTGAAGAACGAGATCAGC-3’, R: 5’-GTAGCCGTGATGCCATTGTCA-3’) was used to detect mutant or K/O mouse Smn1 locus. A primer pair (F: 5’-AAGTGAGAACTCCAGGTCTCCTG-3’, R: 5’-TTCACCGTGTTAGCCAGGATGGTC-3’) was used to detect the human SMN2 gene.

### Preparation of single-cell suspension

Whole spinal cords were obtained from transgenic mice, and the spinal cords were stripped according to established methods and placed in ice-cold phosphate-buffered saline (PBS) [100]. The tissue was minced and dissociated with 0.25% trypsin in a centrifuge tube for 30 min at 28 °C, followed by 1 mg/mL collagenase II for 20 min, with gentle mixing every 10 min. The enzymatic reaction was terminated through the addition of 10% fetal bovine serum (FBS). The sample was filtered using a 70 µm strainer to obtain a single-cell suspension.

### scRNA-seq

Using single-cell 3 ‘Library and Gel Bead Kit V3 (10× Genomics, 1000075) and Chromium Single Cell B Chip Kit (10× Genomics, 1000074), the cell suspension (300–600 living cells per microliter determined by Count Star) was loaded onto the Chromium single cell controller (10× Genomics) to generate single-cell gel beads in the emulsion according to the manufacturer’s protocol. In short, single cells were suspended in PBS containing 0.04% bovine serum albumin (BSA). Approximately 6000 cells were added to each channel, and the target cell recovered was estimated to be approximately 3000 cells. Captured cells were lysed, and the released RNAs were barcoded through reverse transcription in individual GEMs. Reverse transcription was performed on an S1000TM Touch Thermal Cycler (Bio Rad) at 53 °C for 45 min followed by 85 °C for 5 min, and incubated at 4 °C. The complementary DNA (cDNA) was generated and then amplified, and the quality was assessed using an Agilent 4200. According to the manufacturer’s instruction, scRNA-seq libraries were constructed using Single Cell 3’ Library and Gel Bead Kit V3. The libraries were finally sequenced using an Illumina Novaseq6000 sequencer with a sequencing depth of at least 100,000 reads per cell with a pair-end 150 bp reading strategy (performed by CapitalBio Technology, Beijing).

### Bulk-seq

The total RNA of spinal cord tissue was extracted through the Trizol method and sent to Vazyme Company (Nanjing, China) for sequencing. Briefly, ribosomal RNA removal was performed first. Using ribosomal-depleted RNA and random hexamers to synthesize one-stranded cDNA, and then we added buffer, deoxynucleoside triphosphates, and enzymes to synthesize two-stranded cDNA. Double-stranded DNA was purified using VAHTS™ DNA Clean Beads, followed by end repair, A-tailing, and the ligation of sequencing adapters. The cDNA strand containing U was degraded by the uracil-DNA glycosylase enzyme, and finally, PCR enrichment was performed; the PCR product was purified with VAHTS^TM^ DNA Clean Beads to obtain the final strand-specific cDNA library. Illumina HiSeq sequencing was performed after the library was qualified for quality control.

### RNA-seq data processing

First, the raw data of bulk-seq was converted into high-quality clean reads. Then, hisat2 [101] was used to align the clean reads to the reference genome of *Mus musculus*. Fragments per kilobase of exon per million mapped fragments (FPKM) expression values of genes were quantified using cufflinks software [102].

For the 10× genomics data, the Cell Ranger Single-Cell toolkit (version 3.0.0) provided by 10× genomics was applied to align reads and generate the gene-cell unique molecular identifier (UMI) matrix for each sample (https://support.10xgenomics.com/single-cell-gene-expression/software/downloads/latest). The read data were mapped using the corresponding *Mus musculus* reference genome. Different samples were merged using the cellranger aggr function. The obtained raw_feature_bc_matrix was loaded, and Seurat R package (v 3.2.2) was further used for downstream analyses [103]. Further quality control was applied to cells based on three metrics step by step, including the total UMI counts, number of detected genes, and proportion of mitochondrial gene counts per cell. Specifically, cells with <200 detected genes and those with a high detection rate (10%) of mitochondrial gene expression were filtered out. To further remove potential doublets in our data, cells with the number of detected genes >4000 were also filtered out. Additionally, we removed a cluster with a high detection rate of the ribosome (>30%). In addition, genes that were detected in <10 cells were filtered out before further analyses. After quality control, data were normalized and scaled using the SCTransform function [104] , and a percentage of ribosomes was regressed out. We removed the batch effect across different individuals through the identification of anchors between individuals and passed these anchors to the IntegrateData function. For visualization, the dimensionality was further reduced using UMAP or t-Distributed Stochastic Neighbor Embedding. To cluster single cells based on their expression profiles, we used an unsupervised graph-based clustering algorithm, Louvain.

### Cell annotation

The marker genes for each cluster were obtained using the FindAllMarkers function in Seurat package [103]. The MAST algorithm was used to calculate statistical significance. Genes that met the following criteria were considered signature genes: (1) adjusted p-value < 0.01 after Bonferroni correction by using all features in the dataset; (2) log fold-change of the average expression > 0.25; and (3) pct.1 > 0.25 (pct.1: the percentage of cells where the feature is detected in the first group); functional enrichment was performed on these genes to obtain cell type enriched pathways. The main cell types were defined using the SingleR package (https://bioconductor.org/packages/devel/bioc/htmL/SingleR.html). Then, we further checked manually to confirm that reported cell type–specific expressed markers were specific for the corresponding clusters based on the following criteria: pct.1 > 0.6 and pct.2 < 0.4.

### Differential expression between groups

For scRNA-seq, DEGs were analyzed using the FindAllMarkers function in the Seurat package. The MAST algorithm was used to calculate statistical significance. Genes that met these criteria were considered DEGs: absolute log fold-change of the average expression ≥ 0.2 and pct > 0.1 (the percentage of cells where the feature is detected in either group). Functional enrichment was performed on these DEGs.

When calculating the difference in gene expression of bulk-seq, the number of reads that fall into each sample was obtained using htseq-count software [105] ; the data were standardized using the estimateSizeFactors function of the DESeq (2012) R package; and p-value and fold change value were calculated for difference comparison by using the nbinomTest function. Genes with a p-value < 0.05 and a fold difference >2 were defined as DEGs by bulk-seq.

### Functional enrichment analysis

KEGG database and GO category database were used for the functional annotation of DEGs. The enrichment analysis of GO categories was performed by using the R clusterProfiler (v3.14.3) package, and the enrichment analysis of the pathways was performed on hypergeometric distribution by using the R phyper function. The GO categories with a false discovery rate < 0.05 were considered significantly enriched. Although pathways with p < 0.05 were regarded as enriched, only those GO categories or pathways containing ≥ 5 DEGs were kept for further analysis.

### Cell-Cell Communication

To study the interactions between cells and to identify the mechanism by which the molecules communicate at a single-cell resolution, the R package CellChat (version 1.1.3) was applied [45]. Two types of interactions of the CellChatDB.mouse database were used: Secreted Signaling and Cell–Cell Contact. CellChat was used on each interaction separately. Using the aggregateNet function in CellChat, the aggregated cell–cell communication network was calculated, and the signaling from each cell group was visualized. Outgoing or incoming signals of certain cell types were recognized using the function netAnalysis_signalingRole_heatmap.

### Cell differentiation trajectory inference

To infer the differentiation of the trajectory of OPCs, we first used monocle (2.14.0) [106] to infer the pseudo time of each cell (method = “ICA”, ordering_genes = marker genes).

### Immunofluorescence

Mouse spinal cord tissue was fixed with 4% paraformaldehyde (Sinopharm Chemical Reagent Co. Ltd) overnight at 4 °C, embedded in Tissue Freezing Medium optimal cutting temperature compound (O.C.T) after dehydration in a gradient of 10%–30% sucrose solution. Then, 6-µm sections were prepared using a microtome (Leica, Wetzlar, Germany). The sections were rehydrated before immunofluorescence staining. For staining, the block was placed in 5% BSA supplemented with 1% Triton X-100 for 30 min at room temperature. Primary antibodies CD31 (AF3628, R&D Systems) and Col3a1 (sc-271249, Santa Cruz) were diluted in a 1:100 ratio with 0.01M PBS. The slides were placed in the cassette at 4 °C overnight. Then, they were diluted with a secondary antibody [Cy3-labeled Donkey Anti-Goat IgG(H+L); Alexa Fluor 488-labeled Goat Anti-Mouse IgG(H+L), Beyotime, Shanghai, China] in a 1:200 ratio with 0.01M PBS and incubated for 2 h at room temperature. They were washed three times with 0.01M PBS for 15 min each. Images were acquired with a fluorescence upright microscope (Zeiss Microscopy, Jena, Germany). The number of positive signals for each antibody in the spinal cord cross-section participated in the statistics.

## Acknowledgements

The authors thank Professor Yimin Hua from Nanjing Normal University for his valuable suggestions for this study; we also thank Lingfang Zhang from Suzhou Lingdian biotechnology Co. Ltd for his help with the bioinformatics analysis.

## Authors’ Contributions

J.S., L.X. and L.W. initiated and designed the study. J.S., J.Q., Q.Y., Q.J., R.Q., X.W., L.W., and L.X. performed the experiments and analyzed data. J.S., J.Q. and L.X. wrote the paper. All authors read and approved the manuscript.

---

**Sup 1.**
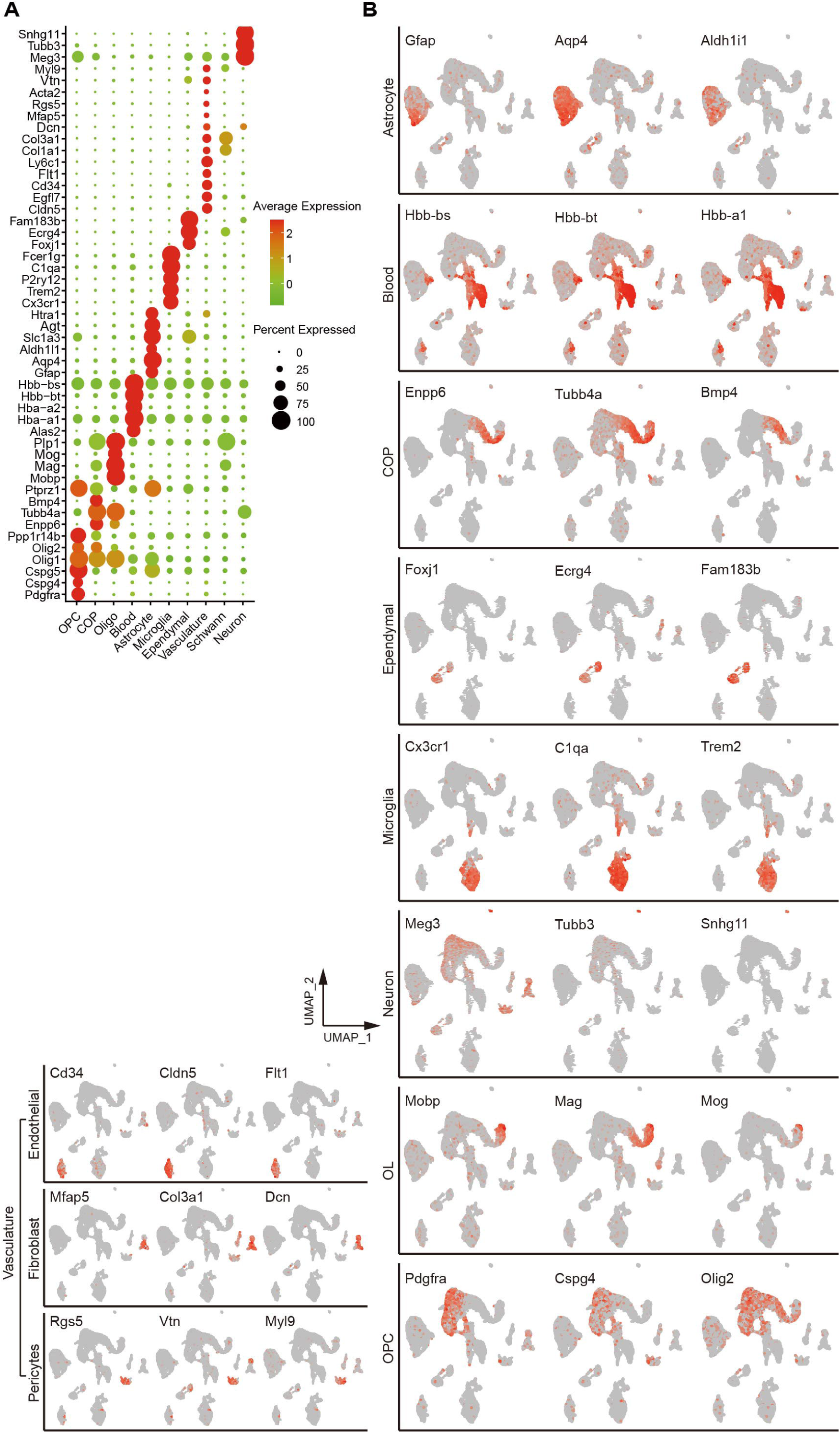
Cell type identification. (A) The average expression of each marker gene in different cell types; the darker the red, the higher the expression. (B) UMAP visualization shows the expression of marker genes in different cell types; the darker the red, the higher the expression.

**Sup 2.**
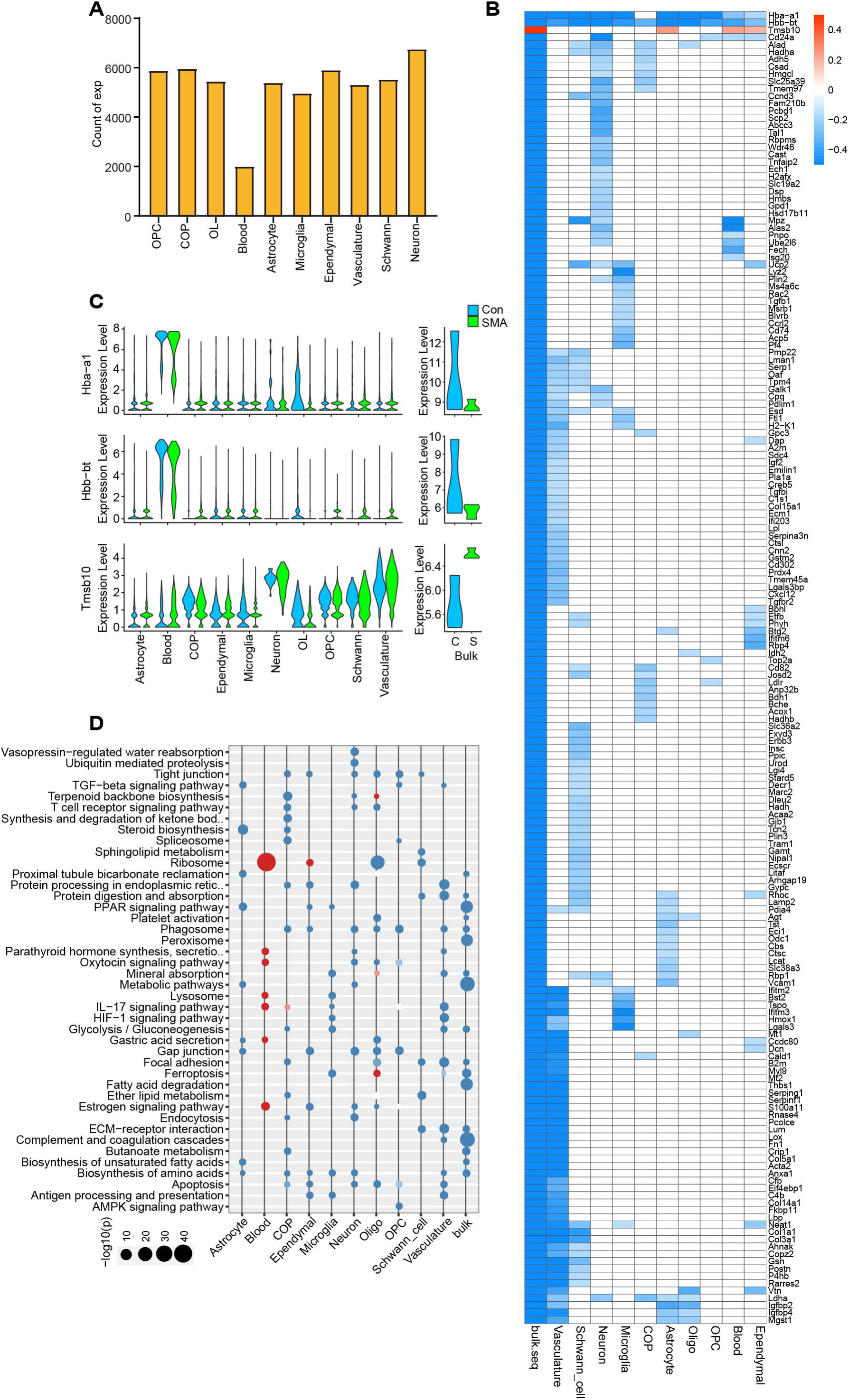
Analysis of DEGs and functions of scRNA-seq and bulk-seq. (A) The number of genes detected through scRNA-seq in each cell type. (B) The cell-type–specific DEGs and bulk-seq DEGs were intersected, and the heatmap shows the expression changes in the intersected genes in different cell types and bulk-seq. (C) Violin plots of Hba-a1, Hbb-bt, and Tmsb10 expression in different cell types and bulk-seq. (D) KEGG pathway analysis of cell-type–specific DEGs and DEGs by bulk-seq; red indicates upregulated pathways, and blue indicates downregulated pathways.

**Sup 3.**
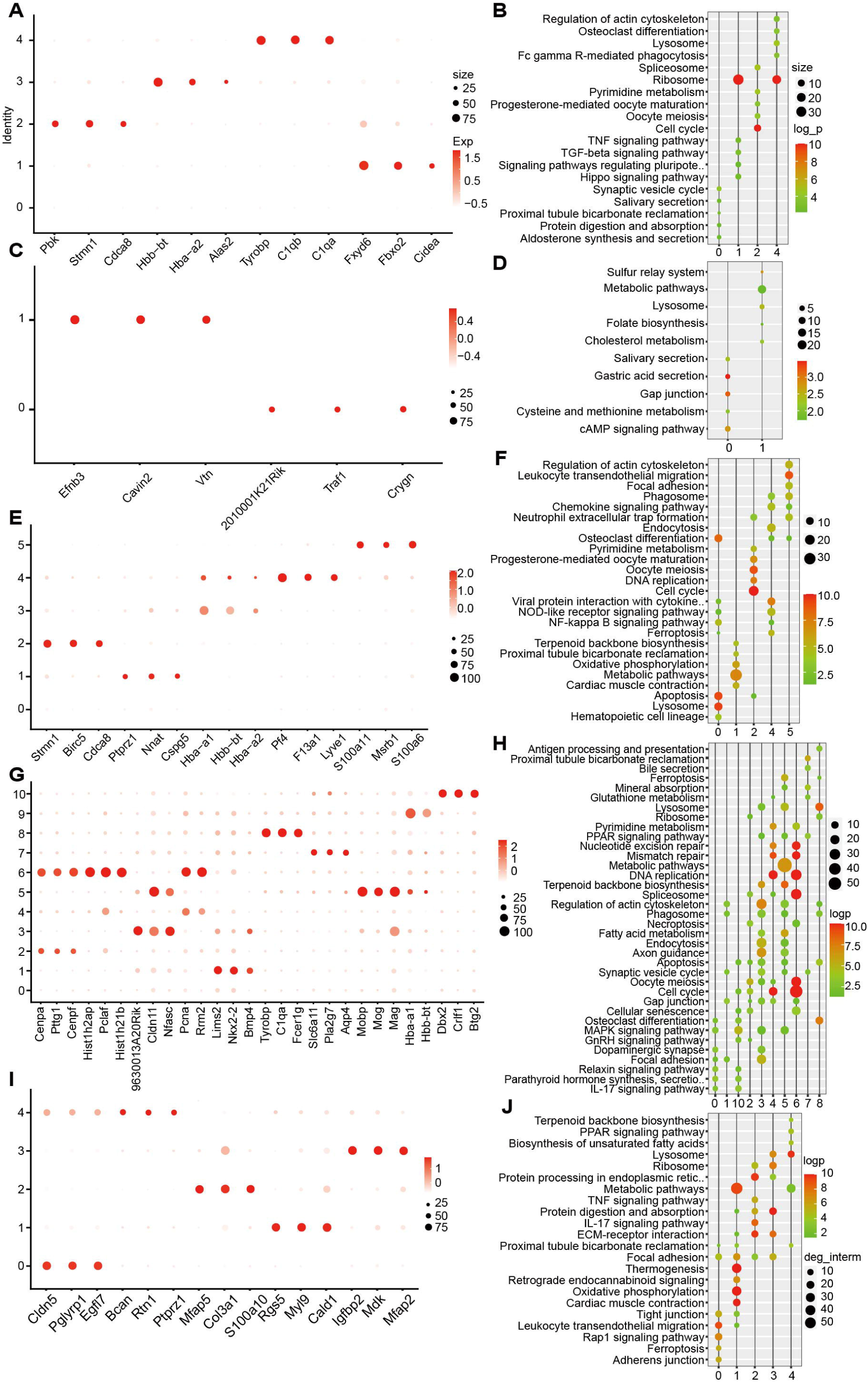
Identify cell subtypes and their DEGs. The average expression of markers of each cell subtype of SMA, the darker the red, the higher the expression; A, C, E, G, and I represent astrocytes, ependymal cells, microglia, OL lineages, and vasculature, respectively. KEGG analysis of marker genes for each cell subtype; B, D, F, H, and J represent astrocytes, ependymal cells, microglia, OL lineages, and vasculature, respectively.

**Sup 4.**
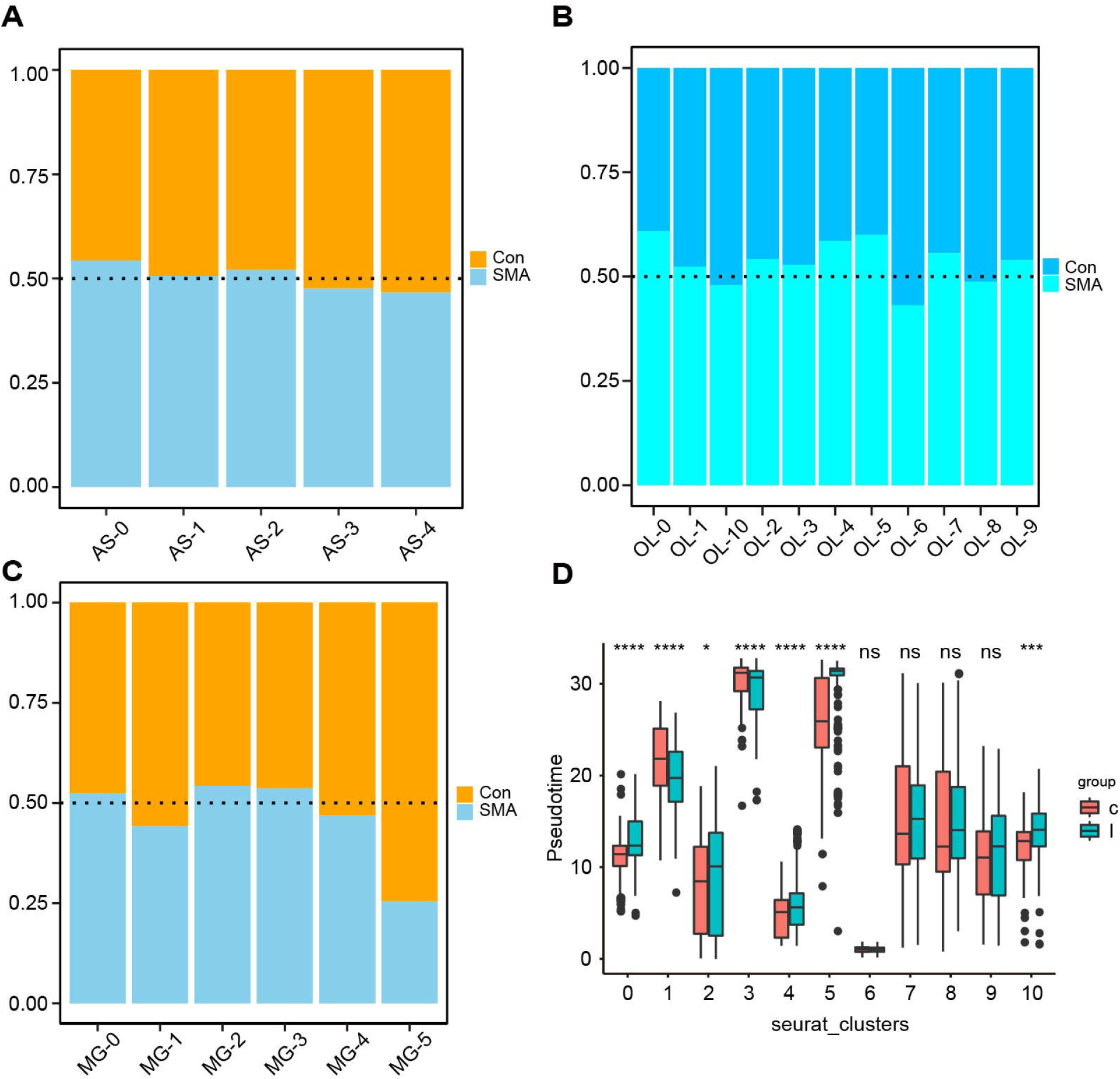
Altered gene expression of glial cell subtypes in the SMA spinal cord. (A–C) Comparison of cell numbers of various subtypes of astrocyte, microglia, and OL lineage in SMA and control mice. (D) Pseudo time analysis of individual cell subtypes in OL lineages. The x-axis represents the cell subtype. The value of the y-axis represents the order of time. The smaller the value, the stronger the characteristics of the progenitor cells. The larger the value, the more mature (later) the cell is. p-values were calculated through the t-test, ns: p > 0.05, *: p < 0.05, ***: p < 0.001, ****: p < 0.0001.

**Sup 5.**
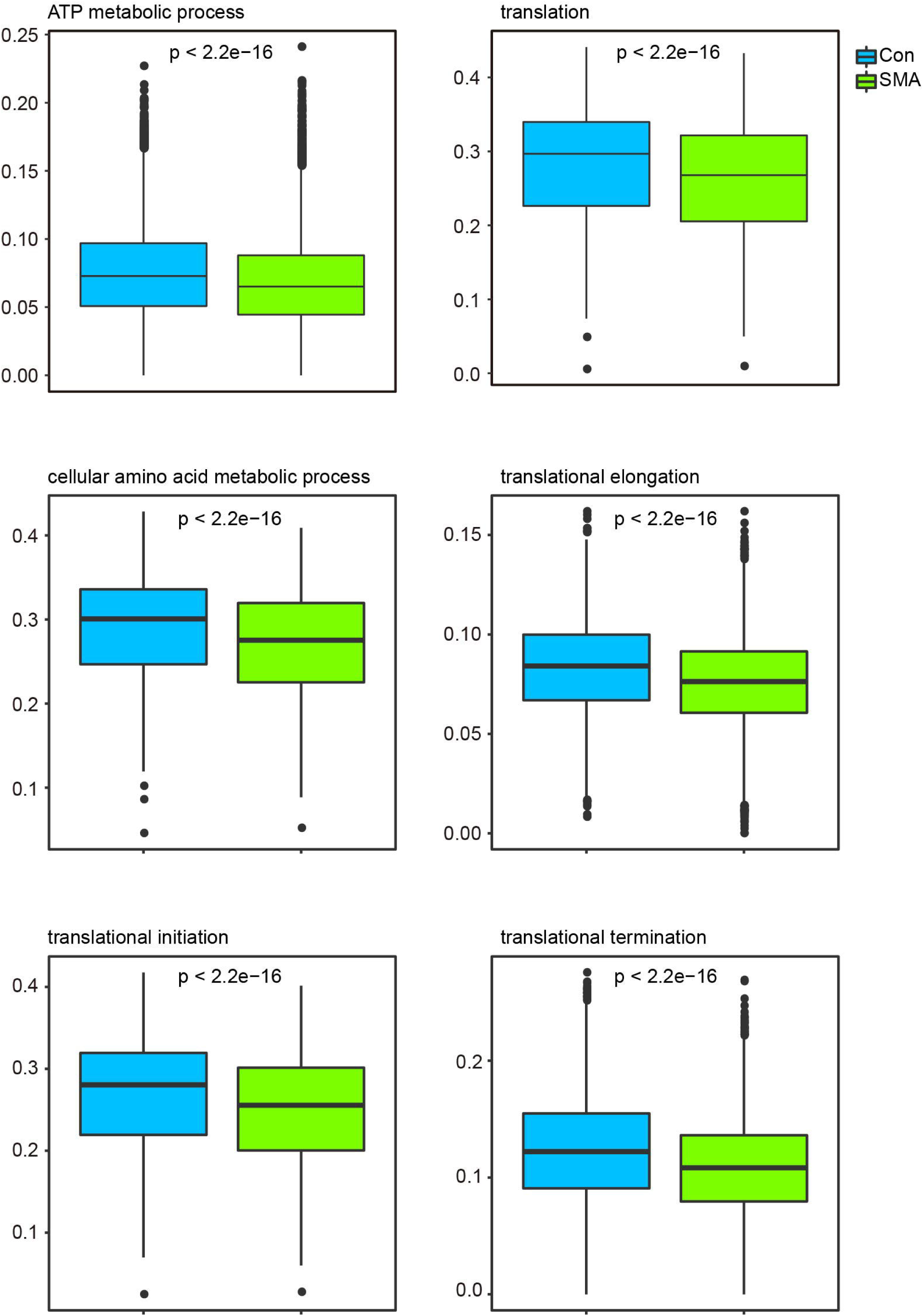
Expression analysis of protein synthesis and energy metabolism-related pathways in SMA and control mice. p-values were calculated by t-test.

Sup File 1. Gene list for bulk-seq and cell-type-specific DEGs. SMA vs. control.

Sup File 2. DEG lists for all cell subtypes. SMA vs. control.

